# Productive foraging sites enhance maternal health and impact offspring fitness in a capital breeding species

**DOI:** 10.1101/2023.10.27.563439

**Authors:** Leila Fouda, Stuart R. B. Negus, Emma C. Lockley, Kirsten Fairweather, Artur Lopes, Anice Lopes, Sandra M. Correia, Albert Taxonera, Gail Schofield, Christophe Eizaguirre

## Abstract

Feeding ecology is an essential component of an organism’s life, but foraging comes with risks and energetic costs. Species in which populations exhibit more than one feeding strategy, such as sea turtles, are good systems for investigating how feeding ecology impacts life-history traits, reproduction and carried over effects across generations. Here, we investigated how the feeding ecology of loggerhead sea turtles (*Caretta caretta*) nesting at the Cabo Verde archipelago correlates with reproductive outputs and offspring fitness. We determined the feeding ecology of female turtles before and during the breeding season from stable isotope analysis of carbon and nitrogen, and correlated isotopic ratio with female and hatchling fitness traits. We found that female turtles feeding at higher trophic positions produced larger clutches. We also found that females with less depleted δ^13^C values, typical of productive foraging areas, had greater fat reserves, were less likely to be infected by leech parasites, and produced heavier offspring. The offspring of infected mothers with less depleted δ^13^C values performed best in crawling and self-righting trials than those of non-infected mothers with less depleted δ^13^C values. Overall, our study shows adult female loggerheads that exploit productive areas build capital reserves that impact their reproductive success and offspring fitness. Together, we uphold the suggestion that not all foraging habitats are equal, and can alter the fitness of populations.

## Introduction

An efficient foraging strategy is one of the most important traits for the survival and Darwinian fitness of an organism (Le Galliard et al., 2004; Stephens & Krebs, 1986). To optimise both foraging and reproductive strategies, some species have evolved long distance migrations between specific habitats (Alerstam et al., 2014; Alerstam & Bäckman, 2018). To undertake these long migrations, individuals must store sufficient resources or be able to acquire them while travelling (Alerstam et al., 2014; Alerstam & Bäckman, 2018). Capital breeders require sufficient energy reserves for both migration and reproduction, unless they successfully complement feeding at the breeding grounds (Wheatley et al., 2008). Therefore, there is a direct connection between the efficiency of foraging and reproductive success, yet this link often remains elusive.

In the marine environment it is challenging to determine feeding ecology directly, where organisms and their prey can be difficult to track (Heerah et al., 2014). Historically, scientists have been limited to direct observations and stomach content analyses (often during necropsy), which provide a snapshot, but possibly limited view, of the feeding ecology of an individual (Hyslop, 1980).To overcome those issues, stable isotope analysis of various tissues (e.g. skin, bone) has become a popular tool, particularly for marine megafauna that are challenging to observe (Crawford et al., 2008; Pauli et al., 2017; Post, 2002). Stable isotope analysis can be performed with minimal impacts on individuals (Guiry et al., 2020; Shiffman et al., 2012) and at large scale (Newsome et al., 2009; Reich et al., 2007; Rubenstein & Hobson, 2004), hence improving our understanding of species feeding ecology by expanding observations from 10’s to 100’s of individuals in a population. For example, stable isotope analysis has unveiled that the pre-breeding diet quality in female Cassin’s auklet (*Ptychoramphus aleuticus*) affected the timing of breeding and egg size, with those exploiting an energetically superior diet breeding earlier and producing larger eggs than their conspecifics (Sorensen et al., 2009).

A property of stable isotope analysis is that different body tissues have different cell turnover rates (Reich et al., 2008; Zanden et al., 2015). Tissues such as teeth and bones have a slower turnover than the skin, which itself has a slower rate than blood plasma (Bearhop et al., 2002, 2004; Reich et al., 2008; Van Klinken, 1999; Vander Zanden et al., 2010). By analysing stable isotope values of different tissues, it is therefore possible to characterise an individual’s feeding ecology across periods of its life cycle. This differential turnover has been leveraged to study southern elephant seals at breeding grounds (*Mirounga leonine*), with vibrissae used to identify the long-term foraging strategies of adult females, while blood plasma was used determine fasting over the breeding and moulting periods (Hückstädt et al., 2012).

An efficient feeding ecology goes beyond an individual meeting its physiological requirements for its own survival, a successful reproduction is also necessary for the persistence of the population and maintaining a high adaptive potential (Clay et al., 2018; Norris et al., 2004; Norris, 2005; Sorensen et al., 2009; Warner et al., 2008; Baltazar-Soares et al., 2014). Parental foraging strategy, diet quality, and energy reserves are directly linked to reproductive output and offspring fitness (Norris et al., 2004; Sorensen et al., 2009; Warner et al., 2008). In the Antarctic fur seals (*Arctocephalus gazella)* for instance, foraging trips by lactating seals correlate with successful breeding and weaning of pups (Gagliano & Mccormick, 2007; Lunn et al., 1994) – directly evidencing the link between diet, food availability and reproduction.

Sea turtles are cryptic marine species that are most accessible when adult females return to their natal rookeries to nest (e.g. Cameron et al 2019). They are capital breeders that can cross entire ocean basins between feeding and breeding grounds, re-migrating every 2 to 4 years depending on the time needed to reach their breeding capital (Bonnet et al., 1998; Plot et al., 2013; Stiebens et al., 2013). Their migration and the subsequent nesting events are energetically costly from the perspectives of breeding, egg production and development, and emerging on the beach to nest (Marn et al., 2017). Loggerhead turtles (*Caretta caretta*) nest on average 3-4 times during the nesting season, and clutch size can vary significantly depending on adult female size and condition (Miller, 1997; van Buskirk & Crowder, 1994).

Loggerhead sea turtles can exhibit dichotomous foraging strategies (Cameron et al., 2019; Hatase et al., 2010; Hawkes et al., 2006). The two main foraging strategies are defined by the habitat used as either oceanic (pelagic) or neritic (coastal). Individuals foraging in the pelagic zone mostly prey upon crustacean and gelatinous prey items in the water column (Frick et al., 2009; Hatase et al., 2010). Neritic foragers, on the other hand, use more productive coastal environments and prey on sessile and slow-moving benthic species such as arthropods and gastropods (Frick et al., 2009; Hopkins-Murphy et al., 2003; Plotkin et al., 1993) as well as cnidaria (McClellan et al., 2010; Wallace et al., 2009).

In this study, we used loggerhead turtles nesting at the Cabo Verde archipelago to explore the impacts of foraging ecology on reproductive success. This population of sea turtles is one of the largest nesting aggregations in the world (Taxonera et al., 2022).Most turtles from this population are characterized as oceanic foragers, with some individuals exploiting areas impacted by upwelling events that enrich the baseline food web in nitrogen. A small proportion (<20%) forage along the continental shelf off Sierra Leone (Cameron et al., 2019; Hawkes et al., 2006). Unlike some other turtle aggregations, some adult females from this population are thought to forage during the breeding period. Furthermore, during the nesting season, Cabo Verde loggerhead turtles’ stable isotope values correlate with parasite infection and offspring fitness (Lockley et al., 2020). It is unknown however whether these correlations stem from the breeding capital of turtles built before the breeding season, or from local supplementary feeding and associated infection risks in Cabo Verde waters.

To address this question, we collected skin and blood plasma samples from nesting females. We tested which of the stable isotope values of the two tissue types correlated best with a measure of capital energy (i.e. skinfold to measure body fat), their reproductive output, and their offspring fitness. We hypothesised that if the breeding capital and reproductive fitness were determined in the foraging ground, stable isotope values of skin tissue would correlate better with reproductive output and fitness measures (including infection rate) than those of blood. In contrast, if the breeding capital was less ecologically relevant than the speculated local supplementary feeding, we hypothesized that stable isotope values of plasma would correlate better with fitness measures.

## Methods

### Data collection

#### Sampling nesting females

Sampling took place on the island of Sal in the Cabo Verde archipelago (Figure 1) during the 2018 nesting season (July–October). To quantify long term change in carbon and nitrogen stable isotopes, 3 mm^2^ of non-keratinised skin tissue was collected from the front flippers of 242 nesting females using a sterile single-use scalpel immediately after egg deposition (Cameron et al., 2019; Stiebens et al., 2013). To investigate short-term differences and tissue variation in carbon and nitrogen stable isotope ratios, blood samples of 1–5 ml in volume were collected from the dorsal cervical sinus of 215 females using a 40 mm 21-gauge needle and 5 ml syringe. Blood samples were refrigerated for up to 48 h before being centrifuged. In the field, females were tagged with Passive Integrated Transponder (PIT) tags to enable their identification upon successive nesting events (Stiebens et al., 2013). Notch to notch curved carapace length (CCL, ± 0.1 cm) and width (CCW, ± 0.1 cm) were measured using a tape measure. Body fat was measured (± 0.1 mm) around the latissimus dorsi muscle using a digital calliper (accuracy ± 0.1 mm) as an indication of health. Each individual was also checked for the presence of *Ozobranchus margoi,* a leech parasite known to impact reproductive output of nesting females (Lockley et al., 2020).

**Figure 1:**
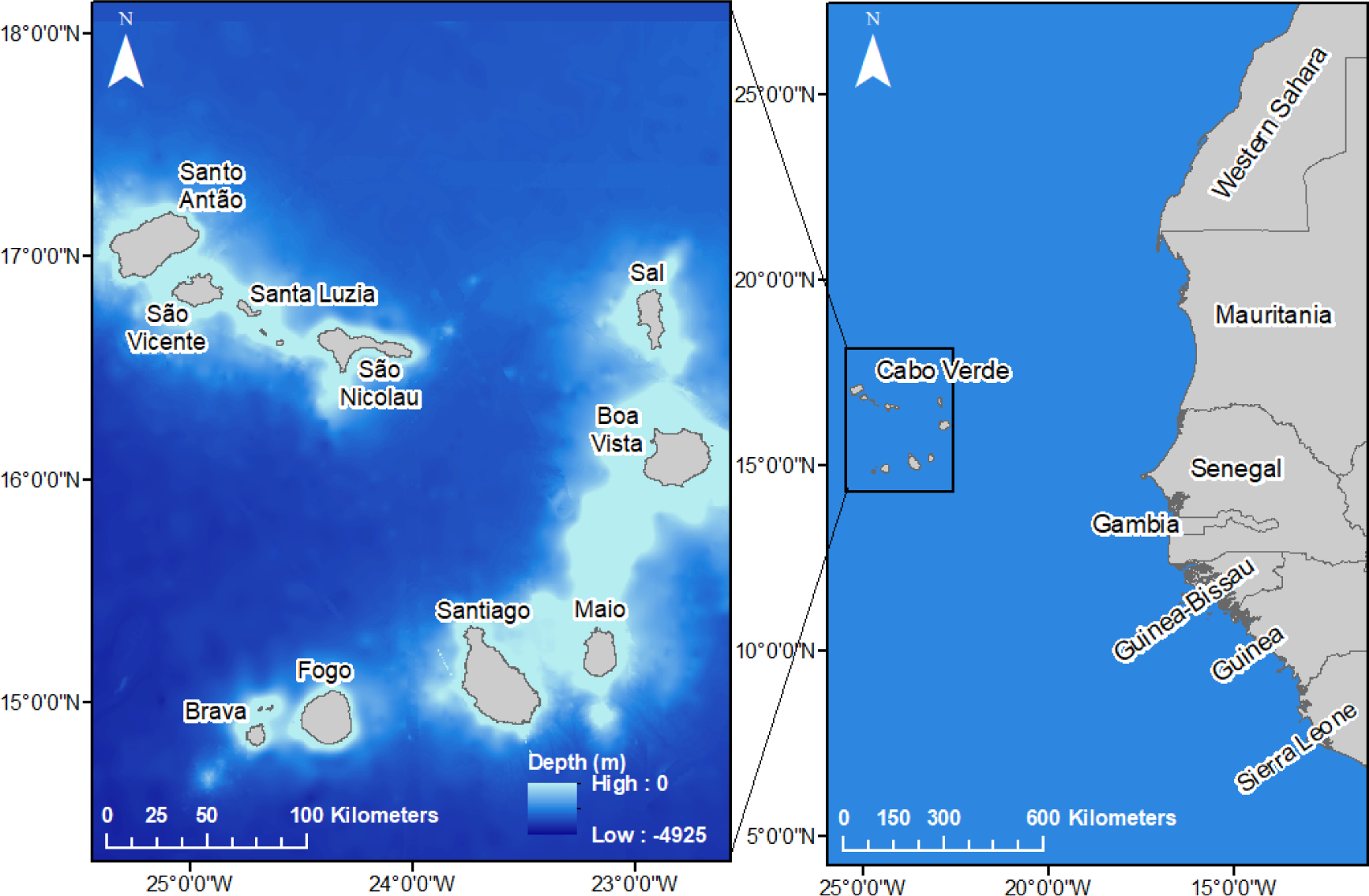
Map of the Cabo Verde archipelago located off the west coast of Africa.

#### Clutch sampling and hatchling fitness measures

To determine reproductive success, the number of eggs in each clutch was recorded and all clutches were relocated to an outdoor hatchery (Lockley et al 2020). After emergence (50–66 days post-oviposition), 20 hatchlings from each nest (N_nest_=88; N_hatchling_=1760) were selected at random for fitness trait measurements. As hatchling size correlates with swimming performance and hatchling dispersal (Scott et al., 2014), hatchlings were measured using digital callipers from notch-to-notch for straight carapace length (SCL, ± 0.01 mm) and weighed (± 0.1 g) so that a Body Condition Index could be calculated:

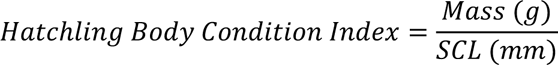

As predator avoidance is critical for hatchling survival, two related traits were also recorded: crawl speed and self-righting time. Crawl speed was measured by recording the time for an individual hatchling to crawl along 50 cm of flat sand, between two pieces of wood, towards a dull red light. Crawl tests were repeated twice and an average was taken (Lockley et al., 2020; Maulany et al., 2012). Self-righting capacity was evaluated by placing a hatchling on its back on a section of flat sand and recording the time taken to right itself (Maulany et al., 2012). Self-righting trials were repeated three times per individual, and if the hatchling took longer than 60 sec to self-right, it was considered to have failed the trial. We measured the number of successful trials (0–3) and the average self-righting time from successful trials in seconds.

### Stable Isotope Analysis

Skin samples were washed in distilled water and dried at 60 °C for 48 h. Between 0.7 μg and 1.3 μg of the sample were placed in tin capsules (4 mm) before being combusted using a continuous flow isotope ratio mass spectrometer (Integra2, Sercon, Crewe, United Kingdom), which simultaneously analysed nitrogen and carbon elements. To verify the accuracy of the readings, an internal standard of casein was run at set intervals (n = 10). Plasma samples were spun before being freeze-dried for 24 h. Between 1.005 μg and 1.156 μg of the dried sample was used. For both tissue samples, the C:N ratio was lower than 3.5, indicating an unlikely bias stemming from lipid enrichment (Post, 2002).

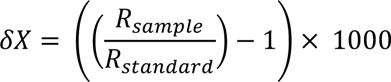

where Χ is ^13^C or ^15^N, R_sample_ gives the ratio of heavy-light isotopes ^13^C/^12^C or ^15^N/^14^N, and R_standard_ gives the ratio of heavy-light isotopes of the standard.

### Statistical analyses

Statistical analyses were conducted in R version 4.0.0 (R Core Team, 2020). All models were backwards selected using Akaike’s information criterion (AIC). In models where fixed variables showed collinearity, we used the residuals of the regression between the two variables to replace one so that both variables remained independent in the models. A generalised linear model was used to regress the body fat measurements against the CCL of the nesting turtles. Residuals were used to create a measure of the adult body fat reserves that accounts for allometric scaling. Linear models were used to determine the relationship between maternal size and body fat reserves. Julian day was included in the model to examine if these relationships changed as the season progressed. Furthermore, due to changes in abiotic factors (like temperature) across the nesting season, we created a variable called “season period” to split the nesting season into three periods of equal duration: Early (09/07/2018 - 12/08/2018), Middle (between 13/08/2018–16/09/2018), and Late (17/09/2018-19/10/2018). This variable was included as an explanatory variable in the models on maternal and hatchling characteristics.

Linear models were used to investigate how stable isotope values changed over the nesting period, as increased δ^15^N is usually considered a starvation signal, while less clear patterns are observed for δ^13^C (Doi et al., 2017). Separate linear models were conducted for δ^15^N and δ^13^C values of skin and plasma to look at their patterns over the nesting season and with tissue type.

Maternal health and size are known to correlate with stable isotope values (Eder et al., 2012; Hatase et al., 2010; Lockley et al., 2020). Therefore, we ran separate linear models for both tissue type to determine the relationship between stable isotope values, maternal body fat reserves, size, and infection status (i.e. parasite presence). The three season periods were also included in the models, as were all two-way interactions.

Following our working hypotheses, we tested whether stable isotope values and body fat reserves were correlated with individual clutch size using linear models and conducted separate models for skin and plasma samples. Skin sample models were used to test for the long-term effect associated with forging area (i.e. during the non-breeding period), and plasma sample models were used to test for the short-term effects of possible supplementary feeding at the nesting ground. The linear models included maternal body fat reserves, CCL, parasite presence, and nesting season period (early, middle, late) and all two-way interactions as explanatory variables. Three-way interactions with season period were also included to investigate the fixed predictor interactions as the season progresses.

We used a series of Linear Mixed Effect Models (LMMs) to test whether components of maternal feeding ecology and body fat reserves were correlated with hatchling fitness measurements. In all LMMs, female ID was used as a random effect to account for hatchlings originating from the same nest. As the incubation duration is known to affect hatchling phenotype, it was included as a fixed predictor (Ferguson & Deeming, 1991; Hatase et al., 2018). CCL is known to correlate with clutch size and maternal investment, and so was also included as a fixed predictor. Separate models were run for skin and plasma. Models focusing on hatchling length and mass included maternal δ^15^N and δ^13^C values, body fat reserves, CCL, parasite presence, season period, and incubation duration as explanatory variables. The hatchling mass model also included SCL as an explanatory variable because larger turtles are expected to be heavier. All two-way interactions were used as explanatory variables, as well as the three-way interactions between stable isotope values and incubation duration. This is because of the known effect of incubation duration on phenotypic characteristics. The LMM models for crawl speed and self-righting additionally included hatchling body condition index (Mass/SCL) to account for hatchling weight on performance.

## Results

Blood and skin samples were collected from 215 and 242 adult females, respectively. Hatchlings from 88 nests were assessed for fitness, with 1760 total hatchlings being tested. Turtle body fat reserves were measured for 227 adult females. Among those, parasites were present on 77 (33.9%) of turtles. In Cabo Verde turtles that have less depleted values of δ^13^C in their skin tissue are considered neritic foragers, compared to their more depleted δ^13^C oceanic foraging conspecifics. More enriched values of δ^15^N in the Cabo Verde turtles is indicative of foraging on food items with a more enriched nitrogen baseline and as such foraging at a higher tropic level (Post, 2002).

### Temporal variation in stable isotope values over the nesting season

As different tissue types have different turnover rates, it is important to evaluate how stable isotope ratios vary over the course of the nesting season. When investigating the link between temporal variation and stable isotope values, we found an interaction between tissue type and Julian day for both carbon and nitrogen isotopes. δ^15^N was higher in skin tissue than blood at the start of the season (F_1,449_ = 66.579, p < 0.001), with values in both skin (F_1,237_ = 24.675, p <0.001) and plasma increasing over the nesting season (F_1,210_ = 10.315, p = 0.002). In comparison, δ^13^C was also higher overall in skin samples than in the blood (F_1,449_ = 1425.800, p <0.001), but there was no change in δ^13^C in skin tissue over the nesting season (F_1,237_ = 0.093, p = 0.76). δ^13^C however did increase over time in plasma samples (F_1,210_ = 22.206, p < 0.001; Figure 2, Supplementary Table S1).

**Figure 2:**
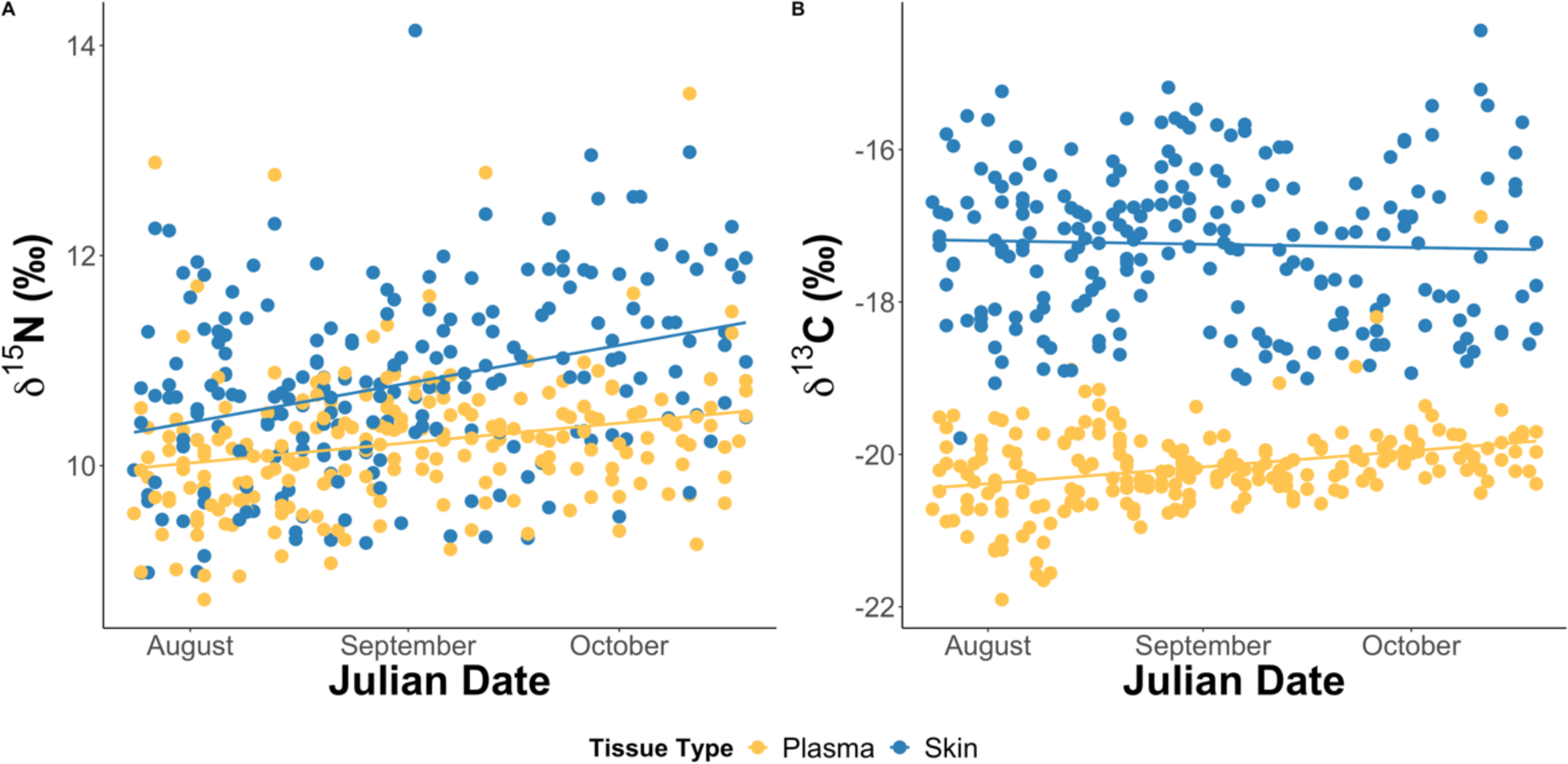
Change in stable isotope values over the 2018 nesting season. A) both skin (n = 242) δ^15^N (F_1,237_ = 24.68, p < 0.001) and plasma (n = 215) δ^15^N (F_1,210_ = 10.32, p = 0.002) increased over the season. B) No trend in skin δ^13^C was seen over time (F_1,237_ = 0.093, p = 0.76), whereas plasma δ^13^C increased (F_1,210_ = 22.21, p < 0.001).

### Maternal Characteristics

Body fat reserve was used as a proxy to evaluate the breeding potential of individual sea turtles. We found that body fat reserves significantly decreased over the nesting season (F_1,225_ = 64.528, p < 0.001). The results remained identical after removal of outliers and showed a significant decrease in the fat surrounding latissimus dorsi muscle of adult females over time (F_1,222_ = 71.522, p < 0.001; Figure 3a). In contrast, the CCL of adult females did not change over the season (F_1,237_ = 2.818, p = 0.095), suggesting that the decline in body fat reserves was not due to a concurrent arrival of a cohort of smaller turtles (Figure 3b).

**Figure 3:**
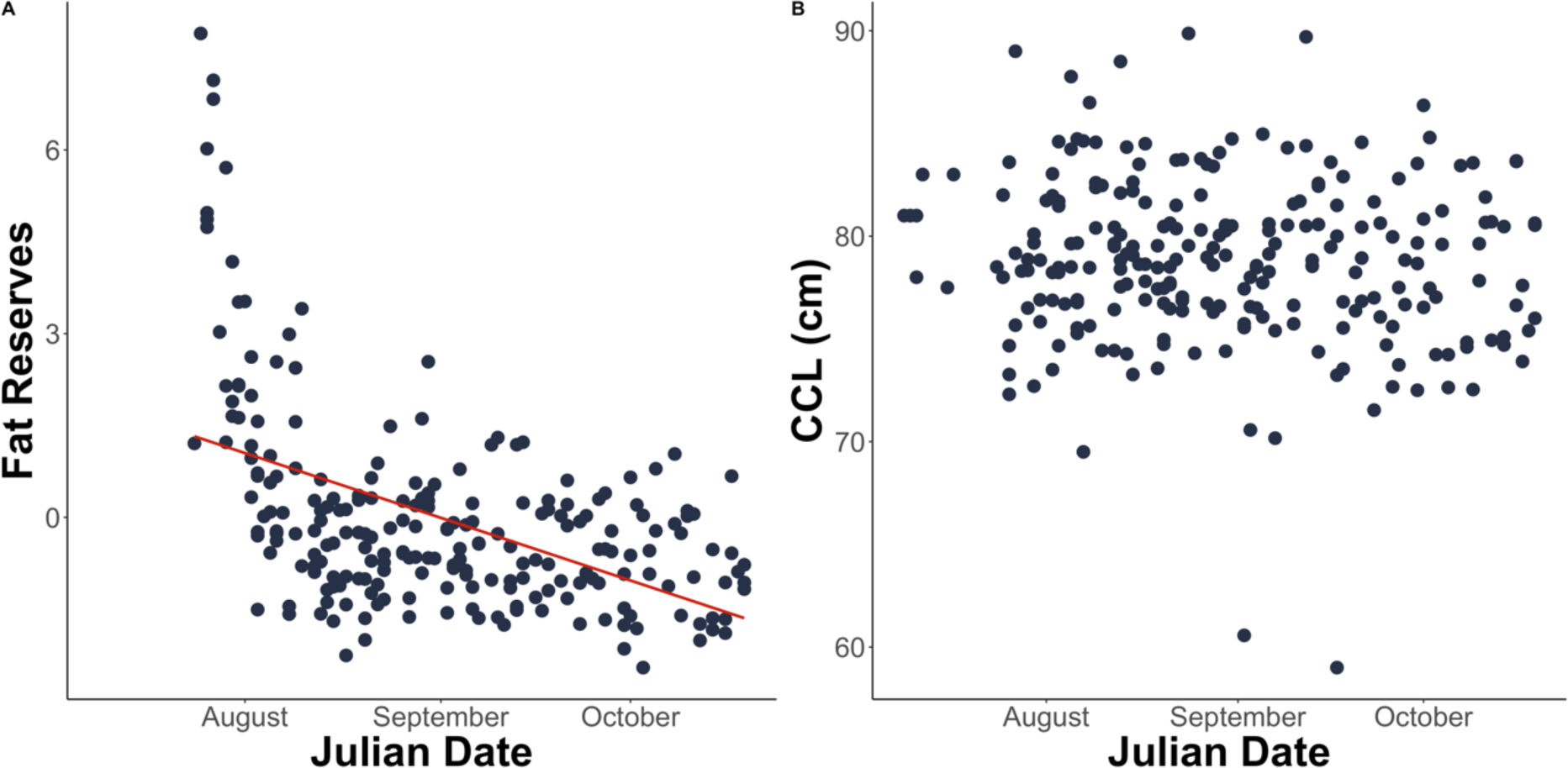
A) Body fat reserves (n = 227) and B) curved carapace length (CCL; n = 242) of adult female loggerhead sea turtle during the 2018 nesting season on Sal Island, Cabo Verde. Body fat reserves of sea turtles significantly declined over the nesting season (F_1,222_ = 71.522 p < 0.001) whereas CCL showed no significant change (F_1,237_ = 2.818, p = 0.095), suggesting no cohort effects in the arrival to the nesting ground.

### Determinants of stable isotope variation in skin and blood plasma

Once we established how body fat reserves vary over the course of the nesting season, we investigated the determinants of isotope variation. Skin δ^13^C ratios positively correlated with female CCL (F_1,207_ = 6.665, p = 0.011; Supplementary Table S2), confirming previously known patterns of larger females feeding at enriched feeding grounds (Cameron et al., 2019).

While at the start of the season, no clear correlation between δ^13^C and body fat reserves were found, this changed as the nesting season progressed, with a positive correlation found in the middle of the season and a negative one at the end (F_2,207_ = 7.695, p <0.001; Figure 4a). These results suggest a very dynamic systems which combine feeding ecology and turtle physiology.

**Figure 4:**
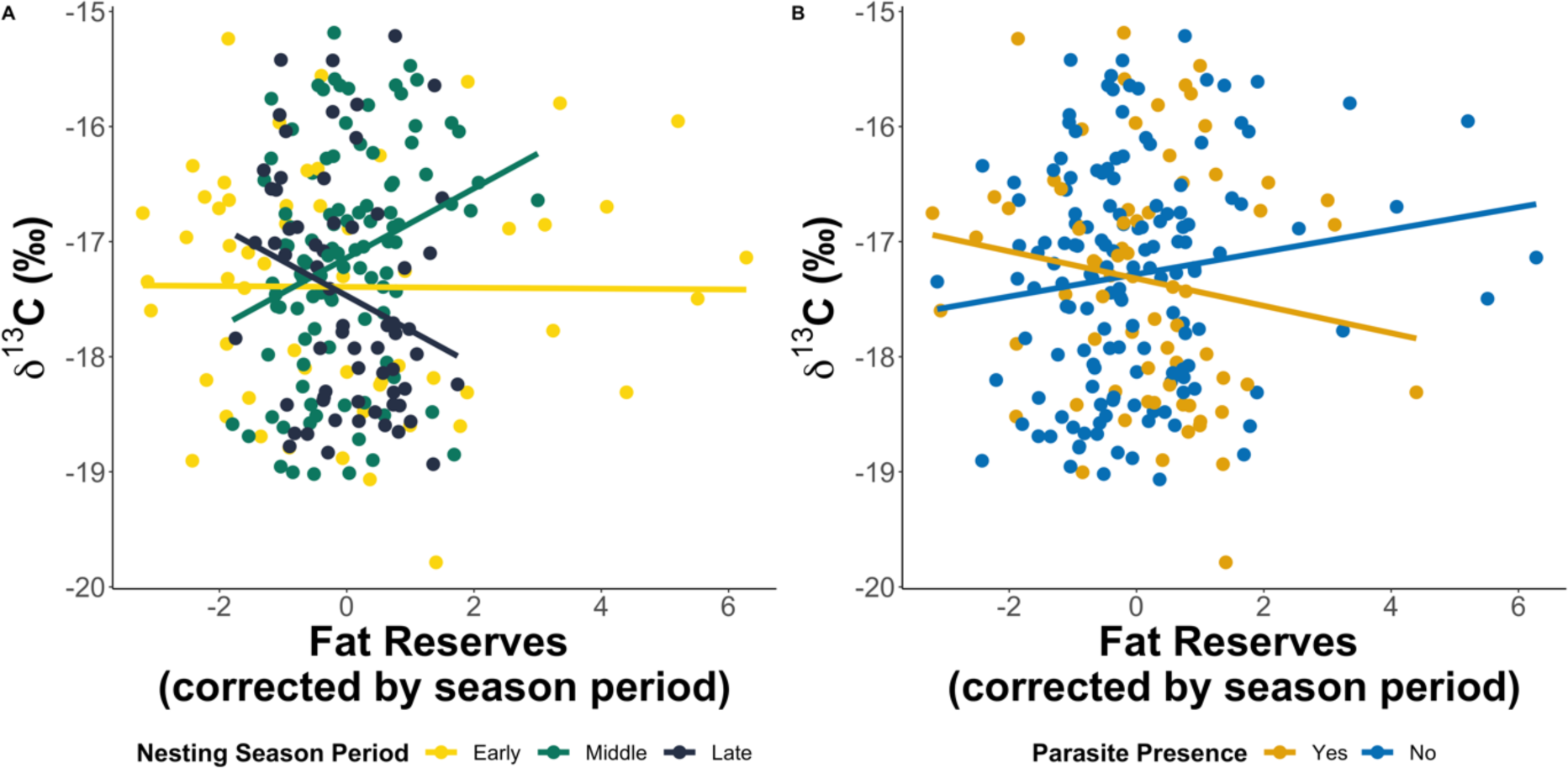
Relationship of turtle (n = 227) skin δ^13^C to body fat (fat reserves regressed against season period) in combination with A) nesting season period (F_2,207_ = 7.695, p <0.001) and B) turtle infection status (F_2,207_ = 5.013, p = 0.026).

Investigating further the effects of turtle physiology, we found that uninfected turtles with high body fat reserves had significantly more enriched δ^13^C values than infected turtles (F_2,207_ = 5.013, p = 0.026; Figure 4b), independent of the nesting season period. This result suggests that oceanic turtles are more likely to be infected by the parasites. This hypothesis is further strengthened by the interaction between CCL and parasite infection on δ^15^N (F_1,208_ = 4.054, p = 0.045), with larger infected turtles having more enriched skin δ^15^N values, suggesting large turtles feed in nitrogen enriched environments but are likely to be exposed to parasites.

When investigating the shorter-term signal of turtle feeding ecology, we first found that plasma δ^13^C correlated negatively with seasonal period (F_2,197_ = 17.715, p < 0.001). The plasma δ^15^N, however, positively correlated with both the period of the season (F_2,196_ = 6.538, p = 0.002) and with an interaction between body fat reserves and CCL (F_1,196_ = 16.125, p < 0.001). As the season progressed, plasma δ^15^N increased, whereby larger turtles with greater body fat reserves showed more enriched δ^15^N. Overall, variation in skin stable isotopes was explained by a combination of seasonal variation and maternal condition, including body fat reserves and infection status. However, differences in plasma isotope ratios were mostly related to seasonal variation.

### Impacts of body fat reserves and foraging on reproductive output

As we found a link between feeding ecology and maternal characteristics, we investigated the link with reproductive output. As expected, we found that larger turtles laid more eggs than their smaller counterparts (N=226, F_1,224_ = 22.489, p < 0.001). Clutch size significantly decreased across the nesting season (F_1,224_ = 31.963, p < 0.001). Interestingly, turtles with more enriched skin δ^13^C values had significantly larger clutches during the early season period (F_2,195_ = 7.536, p < 0.001), but no differences were found during the later season period. The clutch size of turtles was also positively correlated with δ^15^N values in the skin, indicating that turtles foraging at higher trophic levels produce larger clutches (F_1,199_ = 6.010, p = 0.015; Figure 5). Clutch size was not correlated with either plasma δ^15^N or plasma δ^13^C values (δ15N: F_1,187_ = 0.662, p = 0.417; δ13C: F_1,187_ = 0.027, p = 0.866), suggesting the determinant of reproductive output scales with feeding in the foraging habitat.

**Figure 5:**
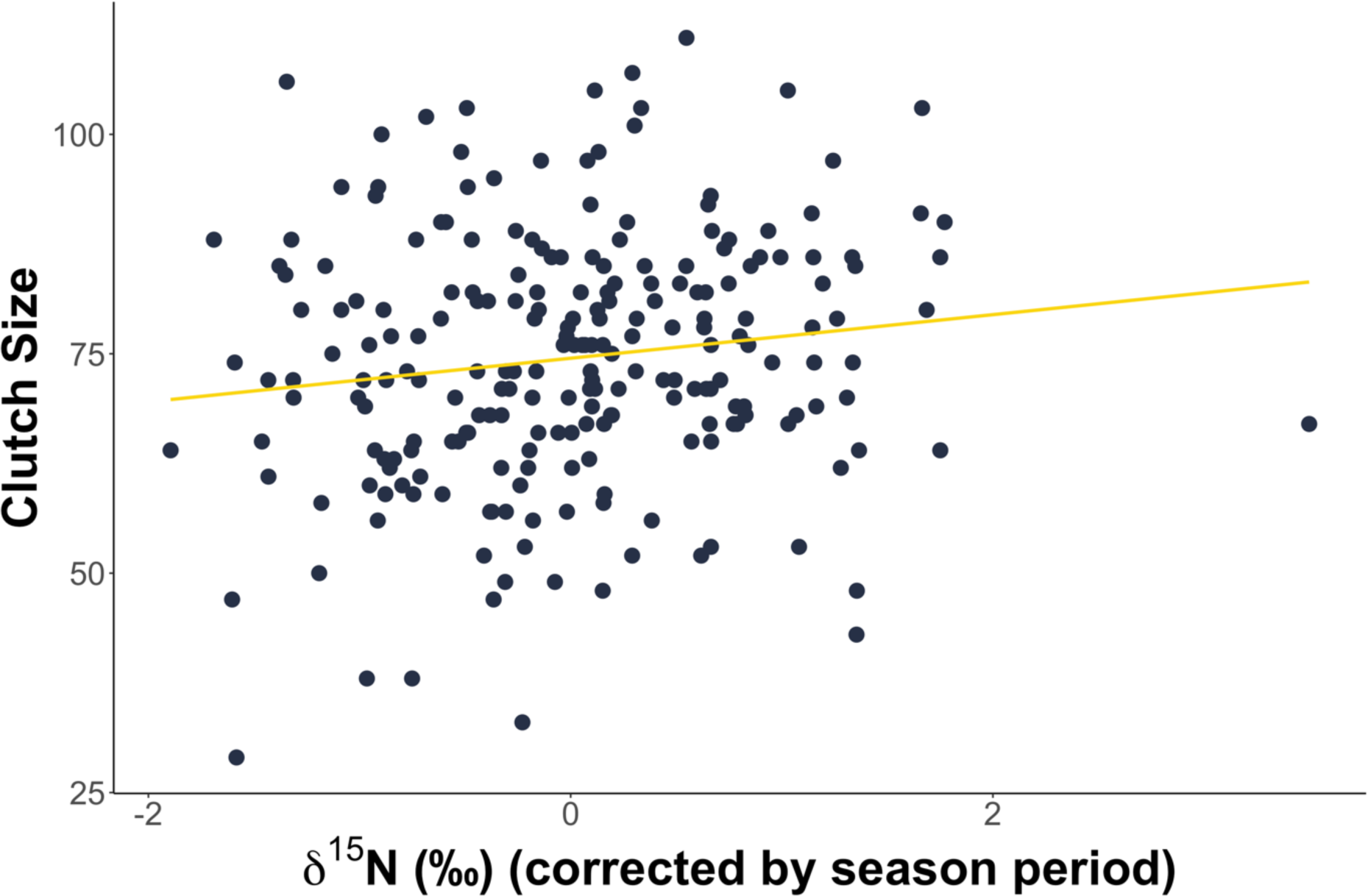
Clutch size (n = 226) positively correlated with δ15N values in the skin (F1,199 = 6.010, p=0.015).

### Effects of maternal condition on offspring fitness

By relocating clutches into an in situ hatchery, we could evaluate the link between maternal feeding ecology, reproductive output all the way to measuring offspring fitness upon emergence. The incubation duration of nests ranged from 50 to 66 days (mean = 54.49, s.d. ± 3.54).

#### Hatchling Size

The length of hatchlings was negatively correlated with maternal body fat reserves (F_1,73_ = 7.29, p = 0.009; Supplementary Table S3) and clutch size (F_2,73_ = 9.97, p = 0.002). As expected in this species, the SCL of hatchlings positively correlated with adult CCL (F_1,73_ = 12.91, p < 0.001). When it comes to the link between maternal feeding ecology and hatchling, we found that the δ^15^N and δ^13^C ratios of skin were not correlated with SCL (δ^15^N: F_1,73_ = 3.61, p = 0.061; δ13C: F_1,73_ = 1.44, p = 0.234). Plasma δ^13^C values however were positively correlated with hatchling SCL (F_1,68_ = 6.98, p = 0.010), unlike plasma δ^15^N (F_1,68_ = 0.42, p = 0.521).

We found that the mothers which foraged in less depleted δ^13^C areas produced hatchlings with a greater mass (F_1,70_ = 7.15, p = 0.009). However, hatchling mass was not significantly correlated with skin δ^15^N (F_1,70_ = 0.12, p = 0.730). While adult females with lower body fat reserves produced significantly heavier offspring (F_1,1586_ = 42.53, p < 0.001); it is important to note that longer offspring were heavier. The offspring of larger mothers were heavier and deposited in bigger clutches than the offspring of smaller mothers (F_1,70_ = 5.42, p = 0.023). The offspring of females with higher plasma δ^13^C were heavier (F_1,1524_ = 11.80, p < 0.001), whereas the offspring of females with enriched plasma δ^15^N were lighter (F_1,1523_=14.19, p < 0.001). However, this relationship was also dependent on offspring SCL. These results suggest that maternal resources are directed into hatchling size and that feeding location in the foraging grounds is important to hatchling size. However, the plasma correlations indicate that foraging habitat alone does not drive hatchling size.

#### Hatchling Performance

Hatchling performance in fitness tests, which is important for hatchling dispersal and survival, was linked to maternal skin δ^13^C. Hatchling crawl speeds and self-righting times were negatively correlated with hatchling body condition index (crawl speed: F_1,1227_ = 6.655, p = 0.018; self-righting: F_1,1190_ = 6.003, p = 0.014). This decrease in speed was expected, as larger individuals tend to be slower. We found that offspring of mothers with parasites and less depleted skin δ^13^C were faster in crawl tests (F_1,69_ = 7.184, p = 0.007) and self-righting trials (F_1,69_=5.622, p=0.021; Figure 6). This suggests hatchling ability to disperse and avoid predation is linked to maternal fitness and foraging habitat.

**Figure 6:**
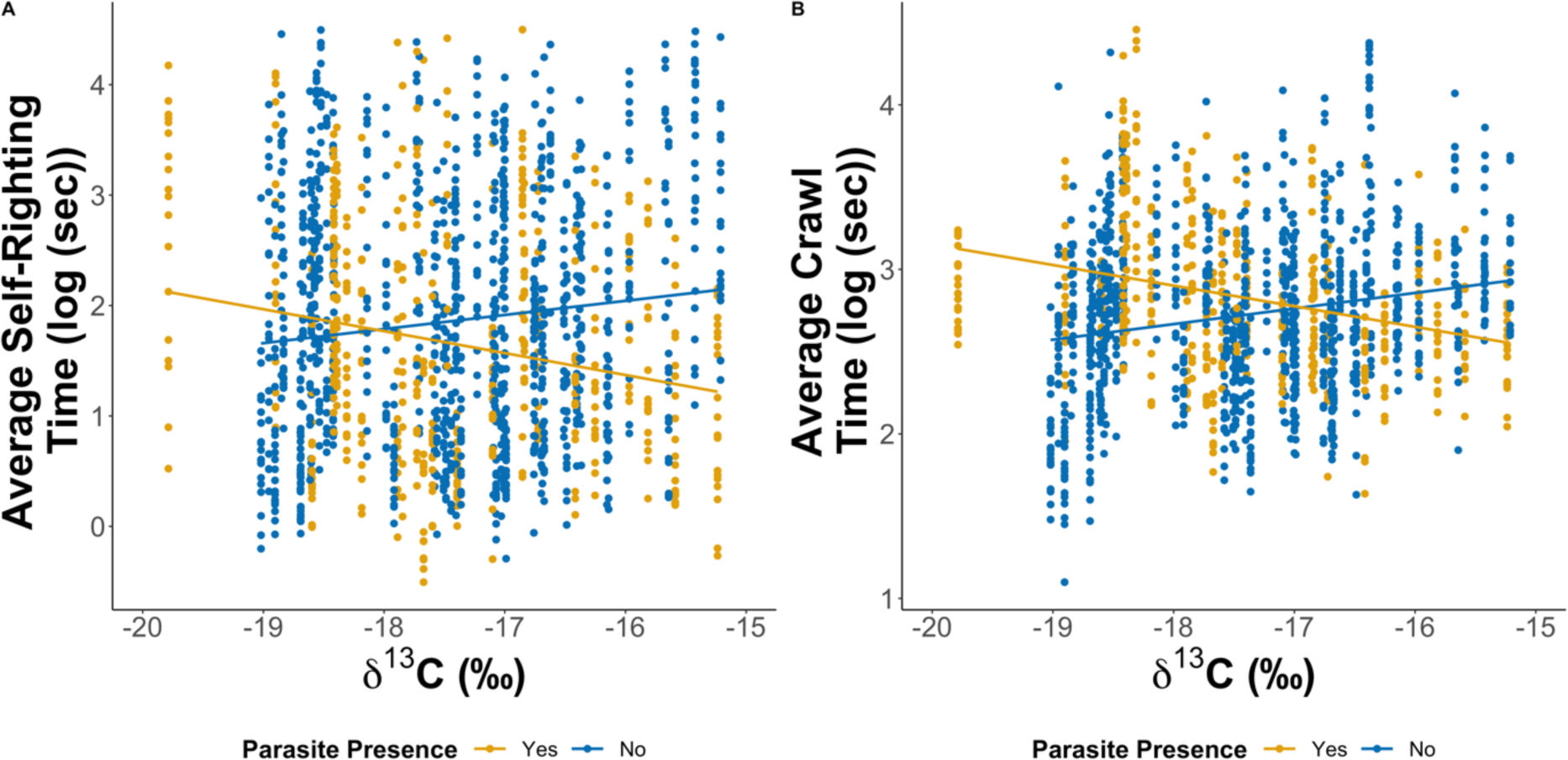
Scatterplots that show hatchling fitness test results with δ^13^C and maternal parasites presence. We found that offspring of infected mothers were faster in A) Self-righting (F_1,69_=5.622, p=0.021), and B) Crawl Speed (F_1,69_=5.734, p=0.019) tests.

Hatchling crawl speeds were correlated with an interaction between nest incubation duration and maternal skin δ^13^C (F_1,69_ = 5.734, p = 0.019). Offspring with a longer incubation duration and from females with lower skin δ^13^C had slower crawl speeds. Nest incubation durations were significantly negatively correlated with self-righting times (F_1,69_ = 6.532, p = 0.013). Additionally, an interaction between female parasite status and clutch size was significantly correlated with self-righting times (F_1,69_=6.747, p = 0.011). While large clutches from females with parasites produced hatchlings with slower self-righting speeds, the opposite was true for uninfected females. Females with higher δ^13^C plasma values had offspring with faster self-righting times (F_1,68_ = 6.42, p = 0.014). Enriched maternal plasma δ^15^N was negatively correlated with self-righting times (F_1,68_ = 7.22, p = 0.009). No significant relationship was found between hatchling crawl times and plasma δ^13^C or δ^15^N (δ^13^C: F_1,66_ = 3.655, p = 0.060; δ^15^N: F_1,65_ = 0.184, p = 0.669). Overall, our results show that hatchling fitness is complex but correlates with maternal stable isotope values, from both maternal blood plasma and skin demonstrating that forces beyond maternal foraging habitat impact hatchling fitness.

## Discussion

Feeding ecology is a critical factor that influences the reproductive output of capital breeders, particularly in migratory species where foraging and breeding grounds can be geographically very distant. In this study, we used stable isotope analysis to test how the feeding ecology and energy stores, in the form of body fat reserves, of adult female loggerhead sea turtles correlates with their reproductive output and the fitness of their offspring. We found that δ^13^C and δ^15^N ratios obtained from skin samples, but not from blood plasma, were the best predictors of nesting females’ fitness. Body fat reserves of nesting turtles decreased as the nesting season progressed, indicating a slow depletion in breeding capital. We found that the healthiest adult females, those uninfected with parasites and with high body fat, had high skin δ^13^C values. Turtles with enriched δ^15^N values, likely foraging in productive coastal oceanic areas, had larger clutch sizes. Together skin δ^15^N and δ^13^C show that not all foraging habitats are equal, and this difference translates into individual fitness and different resource allocation to reproduction with an impact even on offspring. Indeed, females with higher skin δ^13^C values had heavier, and faster offspring. Overall, our findings provide valuable insights into the complex interplay between feeding ecology, and reproductive success, but also reveal the transgenerational carry-over effects of both feeding ecology and health on their offspring fitness.

Firstly, from skin stable isotope analysis, we found that turtles occupy mostly two different niches, linked with oceanic feeding, with one impacted by upwelling events (Cameron et al. 2019). This matches previous studies conducted in Cabo Verde, which identified one more feeding strategy linked to neritic habitat, but only found in turtles nesting on the island of Boa Vista. Those neritic and ocean feeding behaviours have also been confirmed by satellite tracking (Hawkes et al 2008).

Capital breeders store energy in the form of lipid reserves because resource acquisition (e.g. foraging) and resources use (e.g. migration or reproduction) are segregated (Hays, 2000; Kwan, 1994; Perrault et al., 2014). As expected, the body fat reserves of female turtles decreased as the nesting season progressed. Previous studies estimated that turtles weigh roughly 25% more at foraging grounds than at breeding grounds (Hays & Scott, 2013; James et al., 2005). Furthermore, repeated measurements of female green turtles (*Chelonia mydas*) at the nesting grounds on Ascension Island demonstrated that individuals lose 0.22 kg per day over the nesting season (Hays et al., 2002). This weight loss stems from the allocation of capital breeding reserves towards egg production, with no replacement of spent energetic resources.

An important finding in our study was the link between parasite infection, feeding ecology, and energy allocation in female loggerhead turtles. Infected turtles exhibited lower body fat reserves and lower skin δ13C values compared to uninfected turtles. This suggests a trade-off between the energy used to mount an immune response against parasites and food acquisition. Such trade-offs between immune function and resource allocation have been observed in various organisms (Allen & Little, 2011; Sheldon & Verhulst, 1996). Infection with parasites may negatively impact the breeding frequency of females as it requires additional resources to reach the capital breeding threshold while fighting the infection (Allen & Little, 2011). This result confirms a link between feeding ecology and parasite infection in the Cabo Verde loggerhead breeding population, where turtles foraging in relatively poor oceanic environments are more likely to be infected than turtles feeding neritically or in up-welling exposed oceanic areas (Lockley et al., 2020).

Because of infection and unequal quality of feeding grounds, sea turtles could mitigate the allocation of energetic resources to reproduction through supplementary foraging at the breeding grounds. For instance, a mixed foraging strategy has been recorded for lactating Weddle seals (*Leptonychotes weddellii*), that are traditionally capital breeders, but take advantage of occasionally available prey during the reproductive season (Wheatley et al., 2008). Green turtles migrating between their Northern Cyprus breeding grounds and north African foraging grounds appear to use rich coastal waters enroute to replenish resources (Godley et al., 2002). While we did not explicitly identify supplemental feeding in this study, we have observed it directly in this population using on-turtle cameras (BBC, Animal with Cameras 2, personal communication). This supplemental feeding would help top up energy stores lost to energy-intensive breeding and nesting activities.

Our study also revealed significant relationships between feeding ecology, reproductive output. Females with higher skin δ15N values produced larger clutches, indicating either a higher trophic level or baseline-enriched prey items (Hobson, 1999; Post, 2002; Reich et al., 2007; Zbinden et al., 2011). Turtles showing high skin δ^13^C values also produced bigger clutches. This positive correlation between clutch size and high δ^15^N and δ^13^C is explained by turtles foraging at more energy rich areas, such as zones impacted by up-welling events (Cameron et al., 2019), which can provide more resources to allocate to reproduction (Lemos et al., 2020). Similar patterns exist in loggerhead sea turtles nesting at Zakynthos, Greece, with those foraging in the northern Mediterranean (Adriatic and Amvrakikos) showing higher δ^15^N ratios and producing larger clutches than those foraging in the southern Mediterranean (Gulf of Gabes, (Coll et al., 2012; Zbinden et al., 2011)). Those patterns are also seen in marine mammals such as grey seals (*Halichoerus grypus,* (Sparling et al., 2006)).

The benefit of an efficient foraging strategy extends beyond the direct condition of nesting turtles. Specifically, we found that the offspring of female loggerheads with high values of δ^13^C were heavier than those from more δ^13^C depleted mothers. Experimental tests previously showed that hatchlings with longer SCL have better swimming and dispersal abilities (Maulany et al., 2012; Scott et al., 2014). As hatchling mass correlates with length, we can postulate that hatchlings from mothers with high δ^13^C values have better dispersal capabilities (Booth & Evans, 2011; Scott et al., 2014). Infected females with high δ^13^C values produced faster offspring in both the crawl and self-righting trials. A similar result of faster self-righting ability correlating with maternal infection was seen by Lockley et al. (2020). Offspring from females with δ^13^C have better dispersal abilities this clearly demonstrates that some foraging habitats are more beneficial to individual and ultimately population health than others (Scott et al., 2014; Sorci et al., 1994).

Noteworthy, δ13C and δ15N, derived from skin samples proved to be a powerful tool linking feeding ecology and reproduction of sea turtles, unlike stable isotope ratios obtained from blood plasma samples. This discrepancy suggests that plasma samples may serve as a short-term indicator of turtle health, reflecting more recent dietary intake, or lack of, while the determination of clutch size and physiological condition occurs before the nesting season commences. These findings align with previous studies highlighting the differential information provided by skin and plasma samples in stable isotope analysis (Hobson, 1999; Post, 2002; Reich et al., 2007; Zbinden et al., 2011).

Overall, our study showed that individual female turtles that forage in more productive foraging grounds have increased reproductive output and produce fitter offspring. In this capital breeding species, foraging ecology and the capacity to accumulate significant body fat reserves are key to reproductive success, offspring fitness, and could have implications on the survival of the wider population.

## Acknowledgements

We thank all the volunteers and staff from Associação Projeto Biodiversidade, Sal for facilitating this research. LF and CE were supported by the National Environmental Research Council (grant NE/L002485/1, NE/V001469/1). This work was also support by national geographic grants to CE GEFNE69-13, NGS-59158R-19). This work was authorized by the Direção Nacional do Ambiente of Cabo Verde (permit number DNA67/2018).

## Author Contributions

CE and LF conceived the study. LF, CE, EL, SN, AT, KF, ArL, and AnL collected data. AT, KF, SMC, ArL, and AnL facilitated fieldwork and sampling. LF conducted sample processing in the laboratory, the stable isotope analysis, the data analysis. LF and CE interpreted the results. LF drafted the manuscript with the support of CE. GS provided comments and edits on the manuscript.

## Data Availability Statement

Data available from TurtleBase (https://www.qmul.ac.uk/eizaguirrelab/turtle-project/turtlebase/). Code will be made available on GitHub https://github.com/leilafouda/foraging-sites-maternal-health-offspring-fitness

## References

Alerstam, T., & Bäckman, J. (2018). Ecology of animal migration. Current Biology, 28(17), R968–R972. 10.1016/j.cub.2018.04.043

Alerstam, T., Hedenström, A., & Åkesson, S. (2014). Long-Distance Migration: Evolution and Determinants. Oikos, 103(May), 247–260.

Baltazar-Soares, M., Biastoch, A., Harrod, C., Hanel, R., Marohn, L., Prigge, E., Evans, D., Bodles, K., Behrens, E., Böning, C. W., & Eizaguirre, C. (2014). Recruitment Collapse and Population Structure of the European Eel Shaped by Local Ocean Current Dynamics. Current Biology, 24(1), 104–108. 10.1016/j.cub.2013.11.031

Bearhop, S., Adams, C. E., Waldron, S., Fuller, R. A., & Macleod, H. (2004). Determining trophic niche width: A novel approach using stable isotope analysis. Journal of Animal Ecology, 73(5), 1007–1012. 10.1111/j.0021-8790.2004.00861.x

Bearhop, S., Waldron, S., Votier, S. C., & Furness, R. W. (2002). Factors That Influence Assimilation Rates and Fractionation of Nitrogen and Carbon Stable Isotopes in Avian Blood and Feathers. Physiological and Biochemical Zoology, 75(5), 451–458. 10.1086/342800

Bonnet, X., Bradshaw, D., & Shine, R. (1998). Capital versus Income Breeding: An Ectothermic Perspective. Oikos, 83(2), 333. 10.2307/3546846

Booth, D. T., & Evans, A. (2011). Warm Water and Cool Nests Are Best. How Global Warming Might Influence Hatchling Green Turtle Swimming Performance. PLOS ONE, 6(8), e23162. 10.1371/journal.pone.0023162

Cameron, S. J. K., Baltazar-Soares, M., Stiebens, V. A., Reischig, T., Correia, S. M., Harrod, C., & Eizaguirre, C. (2019). Diversity of feeding strategies in loggerhead sea turtles from the Cape Verde archipelago. Marine Biology, 166(10), 130. 10.1007/s00227-019-3571-8

Cherel, Y., Parenteau, C., Bustamante, P., & Bost, C. (2018). Stable isotopes document the winter foraging ecology of king penguins and highlight connectivity between subantarctic and Antarctic ecosystems. Ecology and Evolution, 8(5), 2752–2765. 10.1002/ece3.3883

Clay, T. A., Pearmain, E. J., McGill, R. A. R., Manica, A., & Phillips, R. A. (2018). Age-related variation in non-breeding foraging behaviour and carry-over effects on fitness in an extremely long-lived bird. Functional Ecology, 32(7), 1832–1846. 10.1111/1365-2435.13120

Coll, M., Piroddi, C., Albouy, C., Ben Rais Lasram, F., Cheung, W. W. L., Christensen, V., Karpouzi, V. S., Guilhaumon, F., Mouillot, D., Paleczny, M., Palomares, M. L., Steenbeek, J., Trujillo, P., Watson, R., & Pauly, D. (2012). The Mediterranean Sea under siege: Spatial overlap between marine biodiversity, cumulative threats and marine reserves. Global Ecology and Biogeography, 21(4), 465–480. 10.1111/j.1466-8238.2011.00697.x

Crawford, K., Mcdonald, R. A., & Bearhop, S. (2008). Applications of stable isotope techniques to the ecology of mammals. Mammal Review, 38(1), 87–107. 10.1111/j.1365-2907.2008.00120.x

Eder, E., Ceballos, A., Martins, S., Pérez-García, H., Marín, I., Marco, A., & Cardona, L. (2012). Foraging dichotomy in loggerhead sea turtles Caretta caretta off northwestern Africa. Marine Ecology Progress Series, 470, 113–122. 10.3354/meps10018

Ferguson, M. W., & Deeming, D. C. (1991). Egg incubation: Its effects on embryonic development in birds and reptiles. Cambridge University Press.

Frick, M., Williams, K., Bolten, A., Bjorndal, K., & Martins, H. (2009). Foraging ecology of oceanic-stage loggerhead turtles *Caretta caretta*. Endangered Species Research, 9(2), 91–97. 10.3354/esr00227

Gagliano, M., & Mccormick, M. I. (2007). Maternal condition influences phenotypic selection on offspring. Journal of Animal Ecology, 76(1), 174–182. 10.1111/j.1365-2656.2006.01187.x

Godley, B. J., Richardson, S., Broderick, A. C., Coyne, M. S., Glen, F., & Hays, G. C. (2002). Long-term satellite telemetry of the movements and habitat utilisation by green turtles in the Mediterranean. Ecography, 25(3), 352–362. 10.1034/j.1600-0587.2002.250312.x

Guiry, E., Royle, T. C. A., Matson, R. G., Ward, H., Weir, T., Waber, N., Brown, T. J., Hunt, B. P. V., Price, M. H. H., Finney, B. P., Kaeriyama, M., Qin, Y., Yang, D. Y., & Szpak, P. (2020). Differentiating salmonid migratory ecotypes through stable isotope analysis of collagen: Archaeological and ecological applications. PLOS ONE, 15(4), e0232180. 10.1371/journal.pone.0232180

Hatase, H., Omuta, K., Itou, K., & Komatsu, T. (2018). Effect of maternal foraging habitat on offspring quality in the loggerhead sea turtle (*Caretta caretta*). Ecology and Evolution, 8(6), 3543–3555. 10.1002/ece3.3938

Hatase, H., Omuta, K., & Tsukamoto, K. (2010). Oceanic residents, neritic migrants: A possible mechanism underlying foraging dichotomy in adult female loggerhead turtles (*Caretta caretta*). Marine Biology, 157(6), 1337–1342. 10.1007/s00227-010-1413-9

Hawkes, L. A., Broderick, A. C., Coyne, M. S., Godfrey, M. H., Lopez-Jurado, L.-F., Lopez-Suarez, P., Merino, S. E., Varo-Cruz, N., & Godley, B. J. (2006). Phenotypically Linked Dichotomy in Sea Turtle Foraging Requires Multiple Conservation Approaches. Current Biology, 16(10), 990–995. 10.1016/j.cub.2006.03.063

Hays, G. C. (2000). The Implications of Variable Remigration Intervals for the Assessment of Population Size in Marine Turtles. Journal of Theoretical Biology, 206(2), 221–227. 10.1006/jtbi.2000.2116

Hays, G. C., Broderick, A. C., Glen, F., & Godley, B. J. (2002). Change in body mass associated with long-term fasting in a marine reptile: The case of green turtles (*Chelonia mydas*) at Ascension Island. Canadian Journal of Zoology, 80(7), 1299–1302. 10.1139/z02-110

Hays, G. C., & Scott, R. (2013). Global patterns for upper ceilings on migration distance in sea turtles and comparisons with fish, birds and mammals. Functional Ecology, 27(3), 748–756. 10.1111/1365-2435.12073

Heerah, K., Hindell, M., Guinet, C., & Charrassin, J.-B. (2014). A New Method to Quantify within Dive Foraging Behaviour in Marine Predators. PLOS ONE, 9(6), e99329. 10.1371/journal.pone.0099329

Hopkins-Murphy, S. R., Owens, D. W., Murphy, T. M., Hopkins-Murphy, S. R., & Owens, D. W. (2003). Ecology of immature loggerheads on foraging grounds and adults in internesting habitat in the eastern United States. In A. B. Bolten & B. E. Witherington (Eds.), Loggerhead sea turtles (Vol. 1, pp. 79–92). Washington: Smithsonian Institution Press.

Hückstädt, L., Koch, P., McDonald, B., Goebel, M., Crocker, D., & Costa, D. (2012). Stable isotope analyses reveal individual variability in the trophic ecology of a top marine predator, the southern elephant seal. Oecologia, 169(2), 395–406.

Hyslop, E. J. (1980). Stomach contents analysis-a review of methods and their application. J. Fish Biol, 1741, 1–429. 10.1111/j.1095-8649.1980.tb02775.x

James, M. C., Ottensmeyer, C. A., & Myers, R. A. (2005). Identification of high-use habitat and threats to leatherback sea turtles in northern waters: New directions for conservation. Ecology Letters, 8(2), 195–201. 10.1111/j.1461-0248.2004.00710.x

Kwan, D. (1994). Fat reserves and reproduction in the green turtle, Chelonia mydas. Wildlife Research, 21(3), 257–265. 10.1071/wr9940257

Le Galliard, J.-F., Clobert, J., & Ferrière, R. (2004). Physical performance and Darwinian fitness in lizards. Nature, 432, 502–505. 10.1038/nature03057

Lemos, L. S., Burnett, J. D., Chandler, T. E., Sumich, J. L., & Torres, L. G. (2020). Intra- and inter-annual variation in gray whale body condition on a foraging ground. Ecosphere, 11(4), e03094. 10.1002/ecs2.3094

Lockley, E. C., Fouda, L., Correia, S. M., Taxonera, A., Nash, L. N., Fairweather, K., Reischig, T., Durão, J., Dinis, H., Roque, S. M., Lomba, J. P., dos Passos, L., Cameron, S. J. K., Stiebens, V. A., & Eizaguirre, C. (2020). Long-term survey of sea turtles (*Caretta caretta*) reveals correlations between parasite infection, feeding ecology, reproductive success and population dynamics. Scientific Reports, 10(1), 18569. 10.1038/s41598-020-75498-4

Lunn, N., Boyd, I., & Croxall, J. (1994). Reproductive performance of female Antarctic fur seals: The influence of age, breeding experience, environmental variation and individual quality. Journal of Animal Ecology, 827–840.

Marn, N., Kooijman, S. A. L. M. A. L. M., Jusup, M., Legović, T., & Klanjšček, T. (2017). Inferring physiological energetics of loggerhead turtle (*Caretta caretta*) from existing data using a general metabolic theory. Marine Environmental Research, 126, 14–25.

Maulany, R. I., Booth, D. T., & Baxter, G. S. (2012). The effect of incubation temperature on hatchling quality in the olive ridley turtle, *Lepidochelys olivacea*, from Alas Purwo National Park, East Java, Indonesia: Implications for hatchery management. Marine Biology, 159(12), 2651–2661. 10.1007/s00227-012-2022-6

McClellan, C. M., Braun-McNeill, J., Avens, L., Wallace, B. P., & Read, A. J. (2010). Stable isotopes confirm a foraging dichotomy in juvenile loggerhead sea turtles. Journal of Experimental Marine Biology and Ecology, 387(1–2), 44–51. 10.1016/j.jembe.2010.02.020

Miller, J. D. (1997). Reproduction In Sea Turtles. In L. PL & M. JA (Eds.), The Biology of Sea Turtles, Volume I (pp. 51–81). CRC Press. https://www.taylorfrancis.com/

Newsome, S. D., Tinker, M. T., Monson, D. H., Oftedal, O. T., Ralls, K., Staedler, M. M., Fogel, M. L., & Estes, J. A. (2009). Using stable isotopes to investigate individual diet specialization in California sea otters (*Enhydra lutris nereis*). Ecology, 90(4), 961–974. 10.1890/07-1812.1

Norris, D. R. (2005). Carry-Over Effects and Habitat Quality in Migratory Populations. Oikos, 109(1), 178–186.

Norris, D. R., Marra, P. P., Kyser, T. K., Sherry, T. W., & Ratcliffe, L. M. (2004). Tropical winter habitat limits reproductive success on the temperate breeding grounds in a migratory bird. Proceedings of the Royal Society of London. Series B: Biological Sciences, 271(1534), 59–64. 10.1098/rspb.2003.2569

Pauli, J. N., Newsome, S. D., Cook, J. A., Harrod, C., Steffan, S. A., Baker, C. J. O., Ben-David, M., Bloom, D., Bowen, G. J., Cerling, T. E., Cicero, C., Cook, C., Dohm, M., Dharampal, P. S., Graves, G., Gropp, R., Hobson, K. A., Jordan, C., MacFadden, B., … Hayden, B. (2017). Opinion: Why we need a centralized repository for isotopic data. Proceedings of the National Academy of Sciences, 114(12), 2997–3001. 10.1073/pnas.1701742114

Perrault, J. R., Wyneken, J., Page-Karjian, A., Merrill, A., & Miller, D. L. (2014). Seasonal trends in nesting leatherback turtle (*Dermochelys coriacea*) serum proteins further verify capital breeding hypothesis. Conservation Physiology, 2(1). 10.1093/conphys/cou002

Plot, V., Jenkins, T., Robin, J.-P., Fossette, S., & Georges, J.-Y. (2013). Leatherback Turtles Are Capital Breeders: Morphometric and Physiological Evidence from Longitudinal Monitoring. Physiological and Biochemical Zoology, 86(4), 385–397. 10.1086/671127

Plotkin, P. T., Wicksten, M. K., & Amos, A. F. (1993). Feeding ecology of the loggerhead sea turtle Caretta caretta in the Northwestern Gulf of Mexico. Marine Biology, 115(1), 1–5. 10.1007/BF00349379

Post, D. M. (2002). Using Stable Isotopes to Estimate Trophic Position: Models, Methods, and Assumptions. Ecology, 83(3), 703–718. 10.1890/0012-9658(2002)083[0703:USITET]2.0.CO;2

R Core Team. (2020). R: A language and environment for statistical computing [Computer software]. https://www.R-project.org/

Reich, K. J., Bjorndal, K. A., & Bolten, A. B. (2007). The “lost years” of green turtles: Using stable isotopes to study cryptic lifestages. Biology Letters, 3(6), 712–714. 10.1098/rsbl.2007.0394

Reich, K. J., Bjorndal, K. A., & Martínez del Rio, C. (2008). Effects of growth and tissue type on the kinetics of 13C and 15N incorporation in a rapidly growing ectotherm. Oecologia, 155(4), 651–663. 10.1007/s00442-007-0949-y

Rubenstein, D. R., & Hobson, K. A. (2004). From birds to butterflies: Animal movement patterns and stable isotopes. Trends in Ecology and Evolution, 19(5), 256–263. 10.1016/j.tree.2004.03.017

Scott, R., Biastoch, A., Roder, C., Stiebens, V. A., & Eizaguirre, C. (2014). Nano-tags for neonates and ocean-mediated swimming behaviours linked to rapid dispersal of hatchling sea turtles. Proceedings of the Royal Society B: Biological Sciences, 281(1796), 20141209. 10.1098/rspb.2014.1209

Shiffman, D. S., Gallagher, A. J., Boyle, M. D., Hammerschlag-Peyer, C. M., & Hammerschlag, N. (2012). Stable isotope analysis as a tool for elasmobranch conservation research: A primer for non-specialists. Marine and Freshwater Research, 63(7), 635. 10.1071/MF11235

Sorci, G., Massot, M., & Clobert, J. (1994). Maternal Parasite Load Increases Sprint Speed and Philopatry in Female Offspring of the Common Lizard. The American Naturalist, 144(1), 153–164. 10.1086/285666

Sorensen, M. C., Hipfner, J. M., Kyser, T. K., & Norris, D. R. (2009). Carry-over effects in a Pacific seabird: Stable isotope evidence that pre-breeding diet quality influences reproductive success. Journal of Animal Ecology, 78(2), 460–467. 10.1111/j.1365-2656.2008.01492.x

Sparling, C. E., Speakman, J. R., & Fedak, M. A. (2006). Seasonal variation in the metabolic rate and body composition of female grey seals: Fat conservation prior to high-cost reproduction in a capital breeder? Journal of Comparative Physiology B, 176(6), 505–512. 10.1007/s00360-006-0072-0

Stephens, D. W., & Krebs, J. R. (1986). Foraging Theory. Princeton University Press.

Stiebens, V. A., Merino, S. E., Roder, C., Chain, F. J. J., Lee, P. L. M., & Eizaguirre, C. (2013). Living on the edge: How philopatry maintains adaptive potential. Proceedings of the Royal Society B: Biological Sciences, 280(1763), 20130305. 10.1098/rspb.2013.0305

Taxonera, A., Fairweather, K., Jesus, A., Gonzalves, A., Queiruga, A., Lima, A., Varela-da-Veiga, A., Oujo, C., Lopes, C., Henrique Gomes da Cruz, J., Pina Lomba, J., Patiño, J., E. Medina Suarez, M., Ramos Brás, N., Rendall, P., Correia, S., Roque, S., Eizaguirre, C., Marco, A., … Tiwari, M. (2022). SWOT Report, vol. 17: Cabo Verde: Sea Turtles “In Abundance” (SWOT Report - State of the World’s Sea Turtles, Vol. XVII Vol. 17; SWOT Report). https://www.seaturtlestatus.org/articles/cabo-verde-sea-turtles-in-abundance

van Buskirk, J., & Crowder, L. B. (1994). Life-History Variation in Marine Turtles. Copeia, 1994(1), 66–81. 10.2307/1446672

Van Klinken, G. J. (1999). Bone collagen quality indicators for palaeodietary and radiocarbon measurements. Journal of Archaeological Science, 26(6), 687–695. 10.1006/jasc.1998.0385

Vander Zanden, H. B., Bjorndal, K. A., Reich, K. J., & Bolten, A. B. (2010). Individual specialists in a generalist population: Results from a long-term stable isotope series. Biology Letters, 6(5), 711–714. 10.1098/rsbl.2010.0124

Wallace, B. P., Avens, L., Braun-McNeill, J., & McClellan, C. M. (2009). The diet composition of immature loggerheads: Insights on trophic niche, growth rates, and fisheries interactions. Journal of Experimental Marine Biology and Ecology, 373(1), 50–57. 10.1016/j.jembe.2009.03.006

Warner, D. A., Bonnet, X., Hobson, K. A., & Shine, R. (2008). Lizards Combine Stored Energy and Recently Acquired Nutrients Flexibly to Fuel Reproduction. Journal of Animal Ecology, 77(6), 1242–1249. JSTOR.

Wheatley, K. E., Bradshaw, C. J. A., Harcourt, R. G., & Hindell, M. A. (2008). Feast or famine: Evidence for mixed capital–income breeding strategies in Weddell seals. Oecologia, 155(1), 11–20. 10.1007/s00442-007-0888-7

Zanden, M. J. V., Clayton, M. K., Moody, E. K., Solomon, C. T., & Weidel, B. C. (2015). Stable Isotope Turnover and Half-Life in Animal Tissues: A Literature Synthesis. PLOS ONE, 10(1), e0116182. 10.1371/journal.pone.0116182

Zbinden, J. A., Bearhop, S., Bradshaw, P., Gill, B., Margaritoulis, D., Newton, J., & Godley, B. J. (2011). Migratory dichotomy and associated phenotypic variation in marine turtles revealed by satellite tracking and stable isotope analysis. Marine Ecology Progress Series, 421, 291–302. 10.3354/meps08871

