## Supplementary Table for "Productive foraging sites enhance maternal health and impact offspring fitness in a capital breeding species"

**This file includes:**

Supplementary Tables S1 to S3

Table S1: Summary for linear model results for changes to stable isotope values ($\delta^{13}C$ and $\delta^{15}N$) in the two tissue types (skin and blood plasma) over the 2018 nesting season on Sal. Significant values in bold.

| Variable | d.f. | F | p |
|---|---|---|---|
| **$\delta^{13}C$** | | | |
| Skin~Julian Date | 1, 237 | 0.093 | 0.76 |
| Plasma~Julian Date | 1, 210 | 22.206 | **<0.001** |
| $\delta^{13}C$~Type | 1, 449 | 1424.8 | **<0.001** |
| **$\delta^{15}N$** | | | |
| Skin~Julian Date | 1, 237 | 24.675 | **<0.001** |
| Plasma~Julian Date | 1, 210 | 10.315 | **0.002** |
| $\delta^{15}N$~Type | 1, 449 | 66.579 | **<0.001** |

Table S2: Summary table for best reduced models investigating the correlations of nesting turtle characteristics on stable isotope determinants of $\delta^{13}C$ and $\delta^{15}N$ in the skin and blood samples. Significant values in bold.

| Variable | d.f. | F | p |
|---|---|---|---|
| **Skin $\delta^{13}C$** | | | |
| Fat Reserves | 1, 207 | 0.417 | 0.519 |
| Parasite Presence | 1, 207 | 0.139 | 0.709 |
| CCL | 1, 207 | 6.665 | **0.011** |
| Nesting Season Period | 2, 207 | 3.215 | **0.042** |
| Fat Reserves:Parasite Presence | 1, 207 | 5.013 | **0.026** |
| Fat Reserves:Nesting Season | 2, 207 | 7.695 | **<0.001** |
| **Skin $\delta^{15}N$** | | | |
| Fat Reserves | 1, 208 | 0.484 | 0.488 |
| Parasite Presence | 1, 208 | 0.073 | 0.787 |
| CCL | 1, 208 | 0.17 | 0.681 |
| Nesting Season Period | 2, 208 | 12.265 | **<0.001** |
| Fat Reserves:CCL | 1, 208 | 5.717 | **0.018** |
| Parasite Presence:CCL | 1, 208 | 4.054 | **0.045** |
| Fat Reserves | 1, 197 | 1.031 | 0.311 |
| Parasite Presence | 1, 197 | 0.012 | 0.915 |
| CCL | 1, 197 | 0.222 | 0.638 |
| Nesting Season Period | 2, 197 | 17.715 | **<0.001** |
| Fat Reserves | 1, 196 | 3.528 | 0.062 |
| Parasite Presence | 1, 196 | 3.635 | 0.058 |
| CCL | 1, 196 | 2.165 | 0.143 |
| Nesting Season Period | 2, 196 | 6.538 | **0.002** |
| Fat Reserves:CCL | 1, 196 | 16.125 | **<0.001** |

Table S3: Summary table reporting the best-reduced model testing the effects of maternal feeding ecology and fitness on hatchling health. All models were backwards-selected using AIC. D.F. denotes degrees of freedom. Significant values in bold.

| Variables | d.f. | F | p |
|---|---|---|---|
| **Hatchling Length** | | | |
| Fat Reserves | 1,73 | 7.287 | **0.009** |
| Parasite Presence | 1,73 | 0.083 | 0.774 |
| $\delta^{13}C$ | 1,73 | 3.611 | 0.061 |
| $\delta^{15}N$ | 1,73 | 1.438 | 0.234 |
| CCL | 1,73 | 12.912 | **0.001** |
| Clutch Size | 1,73 | 9.972 | **0.002** |
| Incubation Duration | 1,73 | 1.252 | 0.267 |
| **Hatchling Mass** | | | |
| Fat Reserves | 1,70 | 12.739 | **0.001** |
| Parasite Presence | 1,70 | 0.056 | 0.813 |
| $\delta^{13}C$ | 1,70 | 7.149 | **0.009** |
| $\delta^{15}N$ | 1,70 | 0.12 | 0.73 |
| CCL | 1,71 | 7.702 | **0.007** |
| Clutch Size | 1,71 | 1.114 | 0.295 |
| SCL | 11,595 | 681.202 | **<0.001** |
| Incubation Duration | 1,70 | 2.316 | 0.133 |
| Fat Reserves:SCL | 11,586 | 42.526 | **<0.001** |
| CCL:Clutch Size | 1,70 | 5.413 | **0.023** |
| **Hatchling Crawl Speed** | | | |
| Fat Reserves | 170 | 7.297 | **0.009** |
| Parasite Presence | 1,69 | 7.179 | **0.009** |
| $\delta^{13}C$ | 1,69 | 5.438 | **0.023** |
| $\delta^{15}N$ | 1,69 | 6.34E-05 | 0.994 |

| | | | |
|---|---|---|---|
| CCL | 1,70 | 0.947 | 0.334 |
| Clutch | 1,69 | 3.329 | 0.072 |
| Hatchling Body Mass Index | 11,267 | 5.655 | **0.018** |
| Incubation Duration | 1,69 | 5.405 | **0.023** |
| Fat Reserves: $\delta^{15}N$ | 1,69 | 11.767 | **0.001** |
| Parasite Presence: $\delta^{13}C$ | 1,69 | 7.814 | **0.007** |
| $\delta^{13}C$ :Incubation Duration | 1,69 | 5.734 | **0.019** |
| Clutch Size:Incubation Duration | 1,69 | 3.915 | 0.052 |
| **Hatchling Self-Righting Speed** | | | |
| Fat Reserves | 1,70 | 0.121 | 0.729 |
| Parasite Presence | 1,69 | 6.15 | **0.016** |
| $\delta^{13}C$ | 1,70 | 0.257 | 0.613 |
| $\delta^{15}N$ | 1,69 | 0.145 | 0.705 |
| CCL | 1,72 | 0.093 | 0.761 |
| Clutch | 1,69 | 0.006 | 0.937 |
| Hatchling Body Mass Index | 11,190 | 6.003 | **0.014** |
| Incubation Duration | 1,69 | 6.532 | **0.013** |
| Parasite Presence: $\delta^{13}C$ | 1,69 | 5.622 | **0.021** |
| Parasite Presence:Clutch Size | 1,69 | 6.747 | **0.011** |
